# Genetic Testing Preferences and Intentions in U.S. Patients with Clinically Diagnosed Familial Hypercholesterolemia

**DOI:** 10.1101/587923

**Authors:** Hannah Wand, Amy C. Sturm, Lori Erby, Iris Kindt, William M. P. Klein

## Abstract

**Purpose:** Familial Hypercholesterolemia (FH) is a common Mendelian disorder characterized by elevated LDL cholesterol levels, which if untreated can cause premature heart disease. Less than 10% of cases in the US are diagnosed, and uptake of genetic testing is suboptimal. This study investigates decisionmaking factors associated with intentions to have FH genetic testing among patients clinically diagnosed with FH.

**Methods:** Clinically diagnosed adults with FH and no genetic testing were recruited through the FH Foundation and lipid clinics. Participants completed a survey containing items capturing various reasons to engage in genetic testing.

**Results:** Exploratory factor analysis of survey items identified three factors: (1) aversion to FH genetic information, (2) curiosity regarding medical/family history, (3) and psychological reassurance. Psychological reassurance was, in turn, the only significant predictor of genetic testing intentions. The effect of reassurance was qualified by aversion such that the inverse association between aversion and genetic testing intentions was greater among those with low perceived reassurance.

**Conclusion:** Findings suggest that clinically diagnosed patients’ decisions about FH genetic testing are driven principally by psychological reassurance.

## INTRODUCTION

### Familial Hypercholesterolemia: Current Practices

Familial Hypercholesterolemia (FH) is one of the most common autosomal dominant disorders (prevalence 1/220)^1^ and is characterized by high LDL-C levels resulting in premature cardiovascular disease if left untreated^2^. Early diagnosis and treatment in FH reduces heart disease risk^3^ and with proper and timely management, most individuals live a normal lifespan^3^.

High cholesterol is a common chronic disease with genetic and non-genetic etiologies. Compared to other types of high cholesterol, FH follows a Mendelian inheritance pattern with phenotype present from birth. The underlying genetic etiology of FH is well-described, such that an estimated 60-80% of individuals clinically diagnosed with autosomal dominant FH carry a pathogenic variant in *LDLR, APOB* or *PCSK9*^4^. There has been low awareness and recognition of FH as a Mendelian cause for high cholesterol in the US^5^. FH remains severely underdiagnosed in the US with only up to approximately 10% of affected individuals identified^4^. A diagnosis can be made in the absence of genetic testing (i.e., clinical diagnosis), but a pathogenic variant in one of the genes associated with FH definitively confirms a diagnosis (i.e., genetic diagnosis) and is a tool for family cascade screening.

One potential complication with diagnosis in the US is the lack of standardized diagnostic criteria. Instead, there are several major diagnostic criteria sets (e.g., the Dutch Lipid Clinic Network Score, Simon Broome, AHA and MEDPED) that use various combinations of lipid levels, physical exam, family history, and genetic testing^6^. Due in part to the lack of standardized diagnostic practices, FH can be conflated with high cholesterol that anecdotally “runs in a family” (from, for example, shared lifestyle or multifactorial causes). Recent expert guidelines (released soon after this study was conducted) recommend FH genetic testing and counseling be offered to all patients with suspected FH as part of the diagnostic workup^1^.

Nevertheless, in the US, very few patients diagnosed with FH have had genetic testing^5^. The reasons for the low uptake of genetic testing in this population have not been systematically explored. Some reasons are likely related to healthcare and institutional factors, such as provider attitudes about the value of genetic testing and institutional support for use of this information. In this paper, we consider an equally important set of factors – the perceptions and preferences of the patient.

### Patient Utility of FH Genetic Testing

Genetic testing is ultimately the patient’s decision, yet there are a limited number of studies describing FH patients’ perceived utility of genetic testing. The existing literature generally describes patients’ perceived utility of FH genetic testing *results*, which are not necessarily synonymous with genetic testing *decisions*. These studies seem to highlight two types of benefits to FH genetic testing, psychosocial and health behavior outcomes. In terms of psychosocial benefit, genetic testing can help to reduce uncertainty and stigma, and relieve personal guilt^7,8^. Further, genes are generally not viewed as deterministic and there is usually no sustained emotional distress after receiving genetic information^8,9^. In terms of health behavior, although patients tend to already be engaged in diet and exercise precautions based on elevated perceived risk from family history, a genetic confirmation may help to reinforce these preexisting diet and exercise behaviors^10–13^. Newly identified individuals initiated proper cholesterol-lowering medication(s). Finally, studies have identified utility in using genetic testing for cascade screening in families^14^. One possible limitation of these studies is the nonresponse bias from people declining testing and perhaps a missed opportunity to understand perceived risks or barriers to testing.

It is worth noting that these studies were conducted in countries where the awareness, screening, and diagnosis of FH differs from that in the US, as do the healthcare systems. Awareness about FH in the US is suboptimal, and it is plausible that FH gets conflated with non-genetic and/or non-Mendelian forms of high cholesterol and heart disease. This conflation presents two issues. First, it may present a barrier to genetic testing access and coverage. Second, it may influence patients’ causal attributions about their high cholesterol. For example, if diagnosed with clinical criteria that do not require family history, it is unclear whether patients perceive FH as a Mendelian disorder attributable to a single, “trackable” genetic change. This can influence patients’ perceived utility of genetic testing in terms of management and family screening. In our study, we surveyed participants’ clinical experiences with and knowledge about FH to better understand the context of their decision-making.

We hypothesized that participants’ perceived benefits, risks, barriers, and attitudes regarding FH genetic testing would be associated with intentions to engage in FH genetic testing (as well as how these factors might interact). This hypothesis was informed by genetic testing decision-making frameworks that have been validated in other disease contexts^15^.

### Study Objective

The primary goal of this study was to determine what decision-making factors ultimately predict patients’ intentions to have genetic testing for FH, including a consideration of possible interactions among these decision-making factors. We used genetic testing intentions as a proxy for actual test decisions, as it was not within the scope of this study to offer testing to individuals.

## MATERIALS AND METHODS

### Recruitment

Participants were eligible for the study if they were (1) adults (18 years or older) living in the United States who (2) had a clinical diagnosis of FH and (3) had not had genetic testing for FH at the time of the study. Individuals who had genetic testing for disorders other than FH were eligible. Participants were recruited through: (1) advertisements on the FH Foundation Facebook Discussion Forum and online community, (2) posters in lipid clinics at Johns Hopkins Hospital, Mayo Clinic, Geisinger, and Ohio State University Wexner Medical Center, and OhioHealth, (3) posters sent to provider listservs for the National Lipid Association (NLA), cardiovascular Special Interest Group of the National Society of Genetic Counselors (NSGC), and FH Working Group of NSGC, and (4) an in-person patient forum at the 2017 FH Summit in Miami, Florida. These efforts resulted in a sample of 53 individuals. The same survey link was used for all recruitment sources. Surveys were anonymous and individuals who completed the survey received a $15 Amazon gift card.

### Survey Design

We identified published survey items described as perceived benefits (8 items), risks (10 items), barriers (2 items) and attitudes (15 items) related to genetic testing, and items describing genetic testing intention (7 items)^16–22^. Items were adapted to be FH-specific. Five FH Foundation advocates (diagnosed FH patients) piloted the survey for clarity and comprehensiveness and found these items acceptable. The following are example survey items of each category.

### Sample Survey Items (rated from strongly disagree (1) to strongly agree (5))

1. Benefit Item: “If I knew my high cholesterol was genetic, I would experience less stigma.”
2. Risk Item: “I am concerned about the privacy and confidentiality of my results if I have genetic testing for Familial Hypercholesterolemia.”
3. Barrier Item: “Genetic testing for Familial Hypercholesterolemia is too expensive without insurance coverage.”
4. Attitude Item: “Genetic tests should be available for those who want to use them.”
5. Genetic Testing Intention Item: “I would get the genetic test for Familial Hypercholesterolemia.”

To aid in interpretation, the survey included a knowledge survey about FH^23^ as a proxy for assessing how informed participants were about the condition. In addition, there was a follow-up question asking participants to check what additional information they would need, if any, to make their genetic testing decision (for example “cost information”). We also collected demographic (age, gender, race) and selfreported clinical information (age of diagnosis, current medication, history of cardiovascular events, family history). We assessed illness perceptions and causal attributions using the Brief Illness Perception Questionnaire^24^ with permission from the author (Broadbent). Illness perception items are not discussed here as they are tangential to the principal focus of the current paper.

The survey was determined to be exempt from human subjects review by OHSRP (protocol # 17-NHGRI-00055-1). A copy is available on the Open Science Framework https://osf.io/UAWYG/. Permission to use survey must be obtained by corresponding author (Wand).

### Identifying Significant Genetic Testing Decision-Making Factors

We collapsed items related to genetic testing reasons (benefits, risks, barriers, attitudes) into coherent decision-making factors using exploratory factor analysis (EFA). To assess for sufficient variance in an item, we calculated the mean of each item plus one standard deviation. Items with scores >5.5 (given that most scales had an upper limit of 5) were considered to be susceptible to a ceiling effect and subsequently excluded from the factor analysis due to insufficient variance. This method removed the item “I would get genetic testing for Familial Hypercholesterolemia to satisfy my own curiosity.” The remaining 34 items were included in the EFA. EFA accounts for items with reverse coding.

The resulting factors were independent variables in multiple linear regressions (MLR) to (1) predict genetic testing intention and (2) test for interactions among factors. The dependent variable signifying genetic testing intention was the survey item “I would have the genetic test for Familial Hypercholesterolemia.” We controlled for age and gender given that our sample demographic distribution was skewed toward older individuals and women. We did not observe a significant effect of either variable (data not shown). Post-hoc analysis demonstrated that our MLR was sufficiently powered at 88%.

To test for interaction effects among factors, we calculated pairwise interaction terms between the one significant factor and each of the other two factors. (Given the small sample size, we were limited in the number of coefficients we could use and so we could not include all possible pairwise interactions.) This resulted in an MLR with five coefficients. Post-hoc analysis showed sufficient power at 80%.

### Statistical Analysis Package

We used the R statistical analysis program to perform all statistical analyses using the following packages: ggplot2, matrixStats, nFactors, psych, pwr and corrplot.

## RESULTS

### Sample Characteristics

We collected several baseline measures to characterize our participants (Table 1). Of our 53 participants, many were white (83%) and female (60%). The clinical picture was as expected for FH: many participants had heart disease (53%) and some had a history of heart attack (17%). The average age of diagnosis was 26 years. Clinical management appears suboptimal with only half the sample on statins (47%), and the majority had never been offered genetic testing (75%).

**Table 1.** Demographic and clinical information for sample (n=53) shows sample summary of self-reported demographic information and medical history, including previous engagement with genetic testing. Frequency refers to the number of participants who selected the corresponding answer choice, and proportion is out of the total 53 participants.

| **Demographics** | **Frequency** | **Proportion** |
| --- | --- | --- |
| Female | 32 | 60% |
| White/Caucasian | 44 | 83% |
| Age Categories |  |  |
| 18-25 | 7 | 13% |
| 26-35 | 7 | 13% |
| 36-45 | 17 | 32% |
| 46-55 | 11 | 21% |
| 56 and older | 11 | 21% |
| **Clinical History** | **Frequency** | **Proportion** |
| Currently on statins | 25 | 47% |
| Current heart disease | 28 | 53% |
| History of heart attack | 9 | 17% |
| **Genetic Testing History** | **Frequency** | **Proportion** |
| Offered genetic testing | 9 | 25% |
| Requested genetic testing | 1 | 2% |

Most participants seemed informed on FH-related health information (Table 2). The majority knew the correct prevalence, age of onset, and inheritance of FH (64-85%). Only one third (34%) knew the exact risk for heart disease in FH is twenty times greater than general population, but almost all participants knew the risk is greater without knowing the exact number (91%, data not shown).

**Table 2.** Knowledge Scores about Familial Hypercholesterolemia. includes aggregate sample statistics on knowledge survey, where each row represents a question on the survey with correct answers for reference. The column labeled “answered correctly” refers to the number of participants that answered correctly, and proportion is out of the total 53 participants.

| Knowledge Scores |  |  |
| --- | --- | --- |
| Topic | Answered Correctly | Proportion |
| Inheritance risk<br>(Answer: 50%) | 42 | 79% |
| Heart disease risk<br>(Answer: 20 times population risk) | 18 | 34% |
| Population frequency<br>(Answer: 1/250) | 34 | 64% |
| Age of onset<br>(Answer: Birth) | 45 | 85% |
| Average Score | 2.62 out of 4 |  |

### Decision-Making Factors in Genetic Testing Intention

Survey items related to reasons to have or not have genetic testing collapsed into three independent decision-making factors in the factor analysis. The items within each factor seem to thematically represent: aversion to genetic information (“aversion”), curiosity about medical and family history (“medical curiosity”), and psychological reassurance from testing (“reassurance”) (Table 3). The medical curiosity and reassurance factors were correlated with intention (R= 0.674, 0.806 respectively), as well as with each other (R= 0.747). The aversion factor was not correlated with intention. (R= −0.100).

**Table 3.** Factors identified by exploratory factor analysis. lists the three independent factors identified by exploratory factor analysis (EFA) including: aversion, medical curiosity and reassurance. Factors were above the inflection point of an Eigen plot and had an Eigen value >2. Survey items with an SS loading >0.58 were kept within respective factors. Items are shown here exactly as written in the survey, though EFA takes reverse coding into account. The sample mean score and standard deviation of each item is provided. Cronbach’s alpha was greater than or equal to 0.85 for all factors. Each participant received a factor score for each of the three factors by summing intra-factor items after reverse coding.

| **Factor 1. Aversion (Cronbach's alpha=0.87)** | **Mean** | **SD** | **SS loading** |
| --- | --- | --- | --- |
| Genetic testing for Familial Hypercholesterolemia would cause me a lot of anxiety and/or distress. | 2.264 | 1.430 | 0.73 |
| If I knew my high cholesterol was genetic, it would stigmatize me or my family. | 2.377 | 1.559 | 0.64 |
| If I knew my high cholesterol was genetic, it would decrease my self-confidence. | 2.170 | 1.516 | 0.62 |
| I do not need to know or am not interested in knowing if my high cholesterol is genetic. | 2.038 | 1.285 | 0.81 |
| Genetic testing is unnecessary because I can be diagnosed with Familial Hypercholesterolemia through a cholesterol test. | 2.774 | 1.552 | 0.81 |
| My insurance company would cover genetic testing. | 1.736 | 1.767 | 0.6 |
| **Factor 2. Curiosity about Medical and Family History (Cronbach's alpha=0.85)** |  |  |  |
| Genetic testing should be offered to individuals based on their family history. | 4.113 | 1.235 | 0.64 |
| I am curious about my genetic make-up. | 4.396 | 0.840 | 0.66 |
| I would want genetic testing for common diseases that I am at risk for based on my medical history. | 3.830 | 1.236 | 0.74 |
| I would get genetic testing to learn about the risks for my family to have Familial Hypercholesterolemia. | 3.887 | 1.552 | 0.74 |
| I would get genetic testing because it could inform my treatment options. | 3.887 | 1.515 | 0.58 |
| **Factor 3. Reassurance (Cronbach's alpha=0.86)** |  |  |  |
| If I knew my high cholesterol was genetic, it would encourage me to be healthier. | 3.509 | 1.613 | 0.75 |
| If I knew my high cholesterol was genetic, I would feel less blame. | 3.283 | 1.473 | 0.63 |
| If I knew my high cholesterol was genetic, I would experience less stigma. | 3.226 | 1.552 | 0.79 |
| If I knew my high cholesterol was genetic, I would receive better medical care. | 3.509 | 1.625 | 0.78 |

We tested what decision-making factors (aversion, medical curiosity, and reassurance) were predictive of genetic testing intention using multiple linear regression. Despite the inter-correlation between reassurance and medical curiosity, only reassurance was significantly associated with genetic testing intention (β= 0.149, p<0.0001). Medical curiosity (β= 0.020, p=0.513) and aversion (β= −0.050, p=0.216) were not significant predictors.

We then enhanced this model by adding interactions between reassurance and the other two factors. There was a significant interaction between reassurance and aversion (β=0.025, p=0.001). In addition, aversion becomes significant (β= −0.551, p=0.001) while reassurance loses significance (β=-0.333, p=0.090). The coefficients for medical curiosity and the interaction of medical curiosity with reassurance were not significant (p=0.982 and p=0.974 respectively).

The effect of reassurance was qualified by aversion such that an inverse association between aversion and genetic testing intentions was observed only among those with low perceived reassurance (slope= 0.4698 in high reassurance group vs. slope= −1.2232 in low reassurance group) (Figure 1). High and low were defined as one standard deviation above and below the mean.

**Figure 1.**
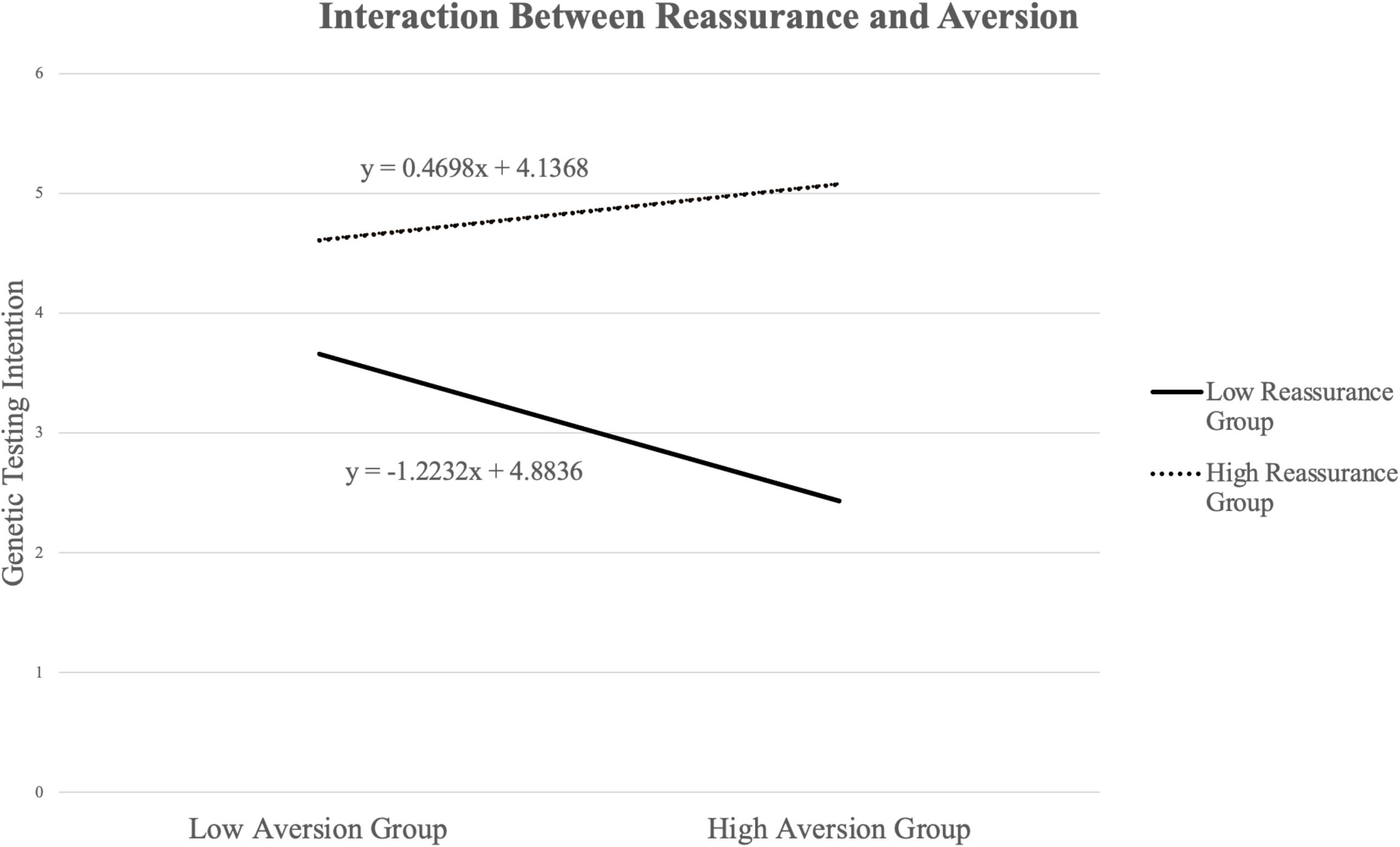
Inverse interaction between reassurance and aversion genetic testing factors. illustrates the interaction relationship between reassurance and aversion. First, we created a linear regression to plot the interaction between aversion and reassurance using aversion and reassurance are the dependent variables, and genetic testing intention is the dependent variable. The resulting equation was (Intention)= 11.779 - 0.539 (aversion) - 0.332 (reassurance) + 0.024 (aversion X reassurance). This equation was used to graph pairwise coordinates for: low aversion & low reassurance, low aversion & high reassurance, high aversion & low reassurance, high aversion & high reassurance. High was defined by one standard deviation about the sample mean and low was defined as one standard deviation below the sample mean. In Figure 1, the y-axis shows intention scores. Colored lines represent low vs. high aversion while the x-axis shows low and high reassurance. A linear line of best fit was used to calculate slope. The low reassurance group has a negative slope of −1.2232, indicating an inverse relationship between aversion and genetic testing, compared to the high reassurance group with a positive slope of 0.4698.

### Requested Genetic Testing Information

At the end of the survey, we asked participants what information they would need about FH genetic testing in order to make their testing decision (Table 4). Participants chose from the list presented in Table 6, with the option to check multiple boxes. The majority of individuals wanted information on insurance and cost. In addition, patients wanted to know how genetic testing would factor into their health management (72-74%). A large proportion of participants also wanted more information about FH in general (40-62%). A minority felt they had all the information they needed (17%), and five participants indicated that they would not need additional information because they would not have testing.

**Table 4.** Requested information on FH genetic testing. includes participants’ options for informational topics they would request in order to make a decision about FH genetic testing. Multiple selections were allowed. Frequency refers to the number of participants who requested that information, with proportion out of the sample of 53 participants.

| **Requested Topics to Inform Testing Decisions** | **Frequency** | **Proportion** |
| --- | --- | --- |
| Cost information | 39 | 74% |
| Insurance coverage information | 38 | 72% |
| Information on the use of genetic information in health management | 33 | 62% |
| Risk assessment based on your personal family history | 29 | 55% |
| Inheritance pattern information | 23 | 43% |
| Disease and symptom information | 21 | 40% |
| None- sufficiently informed to make a decisions | 9 | 17% |
| N/A- would not get testing | 5 | 9% |
| Other | 1 | 2% |

## DISCUSSION

We observed three independent decision-making factors that could be related to FH genetic testing intention: aversion to FH genetic information, curiosity about medical/family history, and psychological reassurance from testing. Although medical curiosity and reassurance were correlated with each other and intention to have testing, only reassurance was significantly predictive of intentions to have genetic testing in the final model. This suggests that patients perceive psychological reassurance as the primary added utility from genetic testing over existing clinical diagnoses. The effect of reassurance was qualified such that, among those with low perceived reassurance, there was an inverse association between aversion and genetic testing intentions. These findings emphasize the importance in the FH pre-test counseling process of discussing potential psychological benefits, in addition to medical benefits, to facilitate decision-making.

These findings are consistent with the existing FH genetic testing patient utility literature, which emphasizes the importance of both psychological and medical perceived benefits of testing. Our study confirms that these two types of benefits separate into independent factors. It has not been previously reported that psychological reassurance is the primary correlate of intention, more so than medical curiosity. This important differentiation of testing benefit suggests that patients prioritize the psychological aspects of FH genetic testing. For providers, the focus on genetic testing utility largely focuses on medical aspects (diagnosis, management, cascade screening) and it is important to know that this may not be the main decision-making factor for patients in our sample.

The complementary idea that barriers related to insurance and privacy concerns would influence testing decisions could not be tested as those items did not coalesce into a separate factor. The top requested informational topics by participants were insurance and cost information. It would be interesting to test whether insurance/cost items collapse into a factor once participants are educated on these topics.

The interpretation and discussion of our study findings should be considered in the context of our sample and recruitment methods. As it was not within the scope of the study to identify and offer FH patients genetic testing, we recruited adults already clinically diagnosed with FH to help ensure that study participants were informed about the condition. As an internal check that participants were informed, we administered a brief FH knowledge survey. The majority of patients knew the answer to all four questions (lowest proportion correct for a question was 64%), assuming we consider awareness about an increased risk of heart disease without knowing the exact risk number to be sufficiently informed. In addition, the average age of diagnosis in our sample (26 years) is roughly two decades before the US average age of diagnosis^25^. Overall, we might characterize this group as informed patients identified through early screening (compared to the general US experience). This is not surprising given our recruitment methods through the FH Foundation and specialized lipid clinics/professional societies that likely overrepresent engaged patients relative to the general FH patient population.

Consequently, the genetic testing predictors and interactions identified by this study may only reflect decision-making in the context of a highly informed and engaged patient sample. These predictors and interactions may not necessarily be representative of the US population or be generalizable to other testing contexts (such as population screening in a healthy population), and results should be replicated in additional cohorts. It is also possible that the items included in our survey were not exhaustive of all the reasons people would choose or decline FH genetic testing. Although we tried to account for this by piloting the survey with five FH advocates (patients) in the FH Foundation, these advocates are also highly informed patients and may not represent reasons that people in the general population would consider pre-test. Lastly, our sample size was relatively small and needs to be replicated with larger cohorts.

It will be important to clarify what patients anticipate in terms of the psychological reassurance gained in testing. Specifically, are patients simply satisfied with the opportunity for psychological reassurance through genetic testing (regardless of testing outcome), or is reassurance tied to the expectation of a specific genetic test result (presumably a positive result to confirm the diagnosis)? If the latter, it will be important to assess for patient understanding about the yield of genetic testing and how a specific result factors into diagnosis during the pre-test counseling process. Pre/post testing studies should evaluate whether patients actually gain a sense of psychological reassurance or reinforcement from genetic testing.

In summary, our findings suggest that genetic testing for FH offers an added dimension of psychological reassurance compared to what a clinical diagnosis provides. This psychological benefit has been reported as a genetic testing benefit in the FH literature, but our study is the first to suggest that it is the significant predictor of genetic testing decisions. Pre-test genetic counseling should assesses for patients’ psychological expectations from genetic testing as there appears to be a significant interaction with aversion to FH genetic information. These testing intention nuances are an informative first step in better understanding patient engagement with precision medicine information, such as FH genetic testing.

## ACKNOWLEDGEMENTS

This work was funded by the National Human Genome Research Institute Intramural Research Training Award as part of the master’s program in genetic counseling at Johns Hopkins/NHGRI. We would like to acknowledge the FH Foundation for piloting and co-sponsoring the study.

## REFERNCES

1. Sturm AC, Knowles JW, Gidding SS, et al. Clinical Genetic Testing for Familial Hypercholesterolemia. J Am Coll Cardiol. 2018;72(6):662–680. doi:10.1016/j.jacc.2018.05.044

2. Nordestgaard BG, Chapman MJ, Humphries SE, et al. Familial hypercholesterolaemia is underdiagnosed and undertreated in the general population: guidance for clinicians to prevent coronary heart disease: Consensus Statement of the European Atherosclerosis Society. Eur Heart J. 2013;34(45):3478–3490. doi:10.1093/eurheartj/eht273

3. Goldberg AC, Hopkins PN, Toth PP, et al. Familial Hypercholesterolemia: Screening, diagnosis and management of pediatric and adult patients. J Clin Lipidol. 2011;5(3):S1–S8. doi:10.1016/j.jacl.2011.04.003

4. Brautbar A, Leary E, Rasmussen K, Wilson DP, Steiner RD, Virani S. Genetics of familial hypercholesterolemia. Curr Atheroscler Rep. 2015;17(4):491. doi:10.1007/s11883-015-0491-z

5. Ahmad ZS, Andersen RL, Andersen LH, et al. US physician practices for diagnosing familial hypercholesterolemia: data from the CASCADE-FH registry. J Clin Lipidol. 2016;10(5):1223–1229. doi:10.1016/j.jacl.2016.07.011

6. Gidding SS, Ann Champagne M, de Ferranti SD, et al. The Agenda for Familial Hypercholesterolemia. Circulation. 2015;132(22):2167–2192. doi:10.1161/CIR.0000000000000297

7. Senior V, Smith JA, Michie S, Marteau TM. Making sense of risk: an interpretative phenomenological analysis of vulnerability to heart disease. J Health Psychol. 2002;7(2):157–168. doi:10.1177/1359105302007002455

8. Weiner K, Durrington PN. Patients & ” Understandings and Experiences of Familial Hypercholesterolemia. Community Genet. 2008;11(5):273–282. doi:10.1159/000121398

9. Collins RE, Wright AJ, Marteau TM. Impact of communicating personalized genetic risk information on perceived control over the risk: A systematic review. Genet Med. 2011;13(4):273–277. doi:10.1097/GIM.0b013e3181f710ca

10. Hagger MS, Hardcastle SJ, Hingley C, Strickland E, Pang J, Watts GF. Predicting Self-Management Behaviors in Familial Hypercholesterolemia Using an Integrated Theoretical Model: the Impact of Beliefs About Illnesses and Beliefs About Behaviors. Int J Behav Med. 2016;23(3):282–294. doi:10.1007/s12529-015-9531-x

11. Senior V, Marteau TM, Weinman J, Genetic Risk Assessment for FH Trial (GRAFT) Study Group. Self-Reported Adherence to Cholesterol-Lowering Medication in Patients with Familial Hypercholesterolaemia: The Role of Illness Perceptions. Cardiovasc Drugs Ther. 2004;18(6):475–481. doi:10.1007/s10557-004-6225-z

12. van Maarle MC, Stouthard MEA, Bonsel GJ. Risk perception of participants in a family-based genetic screening program on familial hypercholesterolemia. Am J Med Genet A. 2003;116A(2): 136–143. doi:10.1002/ajmg.a.10061

13. van Maarle MC, Stouthard MEA, Marang-van de Mheen PJ, Klazinga NS, Bonsel GJ. How disturbing is it to be approached for a genetic cascade screening programme for familial hypercholesterolaemia? Psychological impact and screenees’ views. Community Genet. 2001;4(4):244–252. doi:10.1159/000064200

14. Leren TP, Finborud TH, Manshaus TE, Ose L, Berge KE. Diagnosis of Familial Hypercholesterolemia in General Practice Using Clinical Diagnostic Criteria or Genetic Testing as Part of Cascade Genetic Screening. Public Health Genomics. 2008;11(1):26–35. doi:10.1159/000111637

15. Gooding HC, Organista K, Burack J, Biesecker BB. Genetic susceptibility testing from a stress and coping perspective. Soc Sci Med. 2006;62(8):1880–1890. doi:10.1016/j.socscimed.2005.08.041

16. Delikurt T, Williamson GR, Anastasiadou V, Skirton H. A systematic review of factors that act as barriers to patient referral to genetic services. Eur J Hum Genet. 2015;23(6):739–745. doi:10.1038/ejhg.2014.180

17. Khouzam A, Kwan A, Baxter S, Bernstein JA. Factors Associated with Uptake of Genetics Services for Hypertrophic Cardiomyopathy. J Genet Couns. 2015;24(5):797–809. doi:10.1007/s10897-014-9810-8

18. Meisel SF, Rahman B, Side L, et al. Genetic testing and personalized ovarian cancer screening: a survey of public attitudes. BMC Womens Health. 2016;16(1):46. doi:10.1186/s12905-016-0325-3

19. Schlich-Bakker KJ, ten Kroode HFJ, Wárlám-Rodenhuis CC, van den Bout J, Ausems MGEM. Barriers to participating in genetic counseling and BRCA testing during primary treatment for breast cancer. Genet Med. 2007;9(11):766–777. doi:10.1097GIM.0b013e318159a318

20. van El CG, Cornel MC, ESHG Public and Professional Policy Committee. Genetic testing and common disorders in a public health framework. Eur J Hum Genet. 2011;19(4):377–381. doi:10.1038/ejhg.2010.176

21. Willis AM, Smith SK, Meiser B, Ballinger ML, Thomas DM, Young M-A. Sociodemographic, psychosocial and clinical factors associated with uptake of genetic counselling for hereditary cancer: a systematic review. Clin Genet. 2017;92(2):121–133. doi:10.1111/cge.12868

22. Vermeulen E, Henneman L, van El CG, Cornel MC. Public attitudes towards preventive genomics and personal interest in genetic testing to prevent disease: a survey study. Eur J Public Health. 2014;24(5):768–775. doi:10.1093/eurpub/ckt143

23. Kwok S, Pang J, Adam S, Watts GF, Soran H. An online questionnaire survey of UK general practitioners’ knowledge and management of familial hypercholesterolaemia. BMJ Open. 2016;6(11):e012691. doi:10.1136/bmjopen-2016-012691

24. Broadbent E, Petrie KJ, Main J, Weinman J. The Brief Illness Perception Questionnaire. J Psychosom Res. 2006;60(6):631–637. doi:10.1016/j.jpsychores.2005.10.020

25. deGoma EM, Ahmad ZS, O’Brien EC, et al. Treatment Gaps in Adults With Heterozygous Familial Hypercholesterolemia in the United States: Data From the CASCADE-FH Registry. Circ Cardiovasc Genet. 2016;9(3):240–249. doi:10.1161/CIRCGENETICS.116.001381

